## Supplementary Figs. 1-14 and Supplementary Tab. 1 for "Cortical Reactivation of Non-Spatial and Spatial Memory Representations Coordinate with Hippocampus to Form a Memory Dialogue"

### Author Note

HaoRan Chang https://orcid.org/0000-0001-5981-5181

Majid H. Mohajerani https://orcid.org/0000-0003-0964-2977

Bruce L. McNaughton https://orcid.org/0000-0002-2080-5258

We have no known conflict of interest to disclose.

Correspondence concerning this article should be addressed to HaoRan Chang, Department of Neuroscience, University of Lethbridge, 4401 University Drive, Lethbridge, Alberta T1K 3M4.

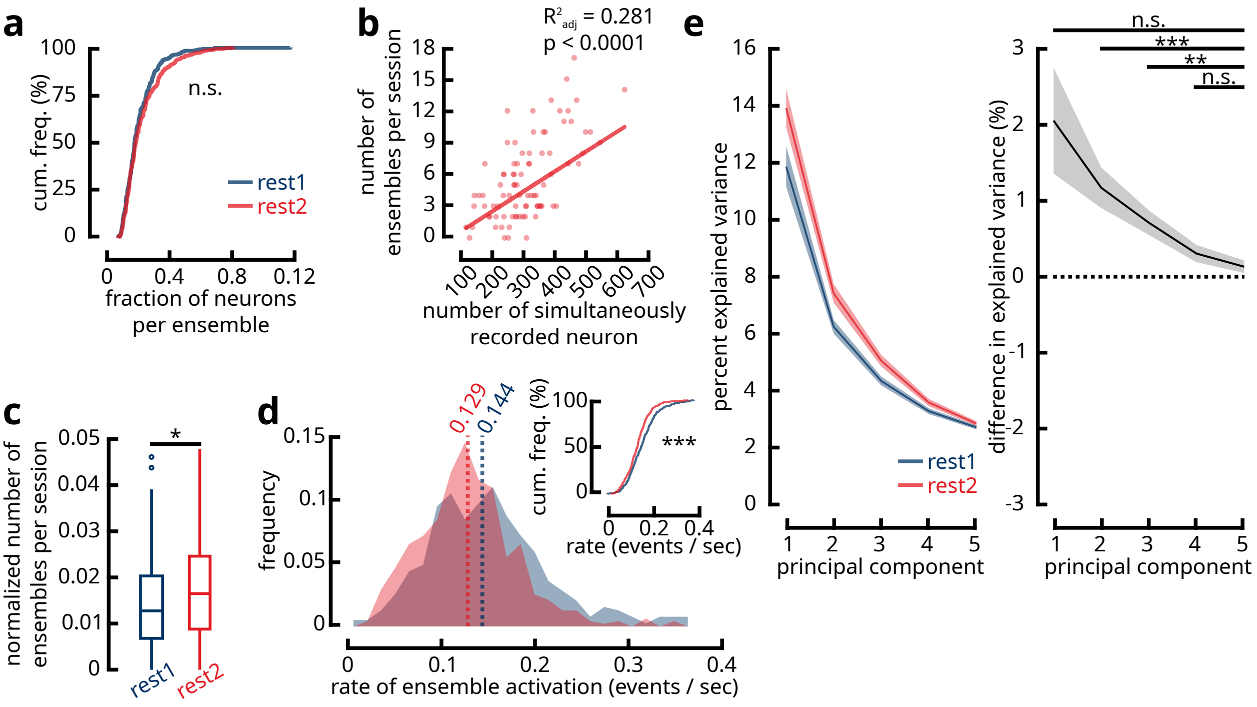

**Supplementary figure 1**

REST2 exhibits more synchronous and sparse population dynamics. (**a**) Cumulative frequency distributions of the fraction of neurons that belong to an ensemble with respect to the total number of simultaneously recorded neurons within a session for REST1 (blue; $n = 86$ sessions; $n = 392$ ensembles) and REST2 (red; $n = 478$ ensembles) (two-sample two-tailed Kolmogorov–Smirnov test; $p = 0.074$). This result suggests that REST1 and REST2 ensembles recruit a similar number of neurons. (**b**) Scatter plot of the number of ensembles per recording session vs. the total number of simultaneously acquired neurons for REST2 ($n = 86$ sessions). Regression analysis revealed a positive correlation between the two quantities. Therefore, as expected, the more neurons are within a recording, the more likely it is to observe ensembles. (**c**) Given that the number of ensembles within a recording is dependent upon the total number of recorded neurons, the former quantity needs to be normalized for appropriate comparisons. The present panel depicts the number of ensembles in REST1 (left) and REST2 (right) expressed as a fraction of the total number of neurons. Paired-samples Wilcoxon signed-rank test revealed that a higher number of synchronous ensembles are found in individual REST2 sessions ($n = 86$ sessions; $p = 0.001$), which suggests that the population activity became more synchronous following active navigation. (**d**) Histogram representation of the average rate of activation (events per second) of REST1 (blue; $n = 392$ ensembles) and REST2 (red; $n = 478$ ensembles) ensembles. The median activation rate of REST1 ensembles was higher than that of

REST2 ensembles (two-tailed Mann-Whitney U-test; $p < 0.001$). The same data is depicted in the form of empirical cumulative distribution functions in the inset (two-sample two-tailed Kolmogorov-Smirnov test; $p < 0.001$). This result indicates that REST2 ensembles expressed higher temporal sparsity in their activities. (**e**) To further verify that REST2 population dynamics show a higher degree of synchrony, PCA analysis was conducted over the correlation matrices of time series vectors of simultaneously recorded neurons in REST1 and REST2 separately. The first five principal components were extracted and the percentage of the total variance explained by each component was scrutinized (**left**: lines show mean±s.e.m.). For these paired samples ($n = 86$ sessions), we took the difference in percentage explained variance between REST2 and REST1 for each component (**right**: line shows mean±s.e.m.) to test whether REST2 components reliably explain more of the variance (one-way repeated measures ANOVA with Greenhouse-Geisser correction; residuals approximately normal; significant difference between components with $p = 0.019$). Post-hoc tests using the fifth component as a baseline reference suggests that the second and third components in REST2 accounted for more of the variance in the population dynamics than their REST1 counterparts (n.s. $p \geq 0.05$; * $p < 0.05$; ** $p < 0.01$; *** $p < 0.001$; p-values were Bonferroni-adjusted). This suggests that a higher degree of correlation existed in REST2 neuronal populations, where more of the variance in the data can be explained by fewer orthogonal vectors.

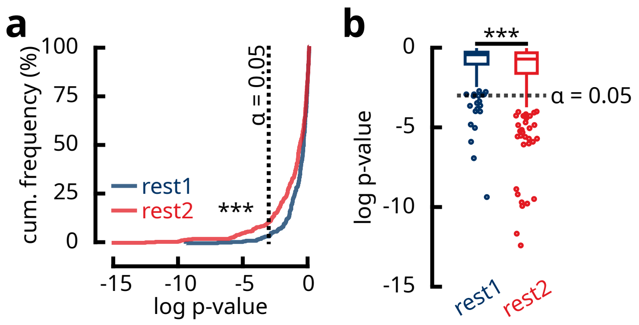

**Supplementary figure 2**

Hypergeometric modelling of the fraction of spatially-selective cells in rest ensembles. For each ensemble ($n = 392$ REST1 ensembles; $n = 478$ REST2 ensembles), the number of spatially-selective cells ($k$) is counted out of the total number of cells that are part of the ensemble ($n$). The total number of spatial cells ($K$) within the corresponding recording session, which contains a total of $N$ simultaneously imaged cells are also counted. A one-tailed Fisher's exact test was conducted on these values. Intuitively, the p-value returned from this test reflects the likelihood of obtaining a certain number of spatial cells in an ensemble of a given size by drawing at chance from the sample population. Small p-values indicate that the fraction of spatial cells within an ensemble is higher than what is expected at chance level. P-values were log-transformed for better detection of low-probability events. **Left**: Empirical cumulative distribution functions of $\log(p_{\text{values}})$ for REST1 and REST2 ensembles. A greater number of REST2 ensembles contained a large fraction of spatially-selective neurons (two-tailed two-sample Kolmogorov-Smirnov test; $p < 0.001$). Dotted line represents significant $\alpha$ level. **Right**: Same as **left**, but represented as boxplots (two-tailed Mann-Whitney U-test; $p < 0.001$).

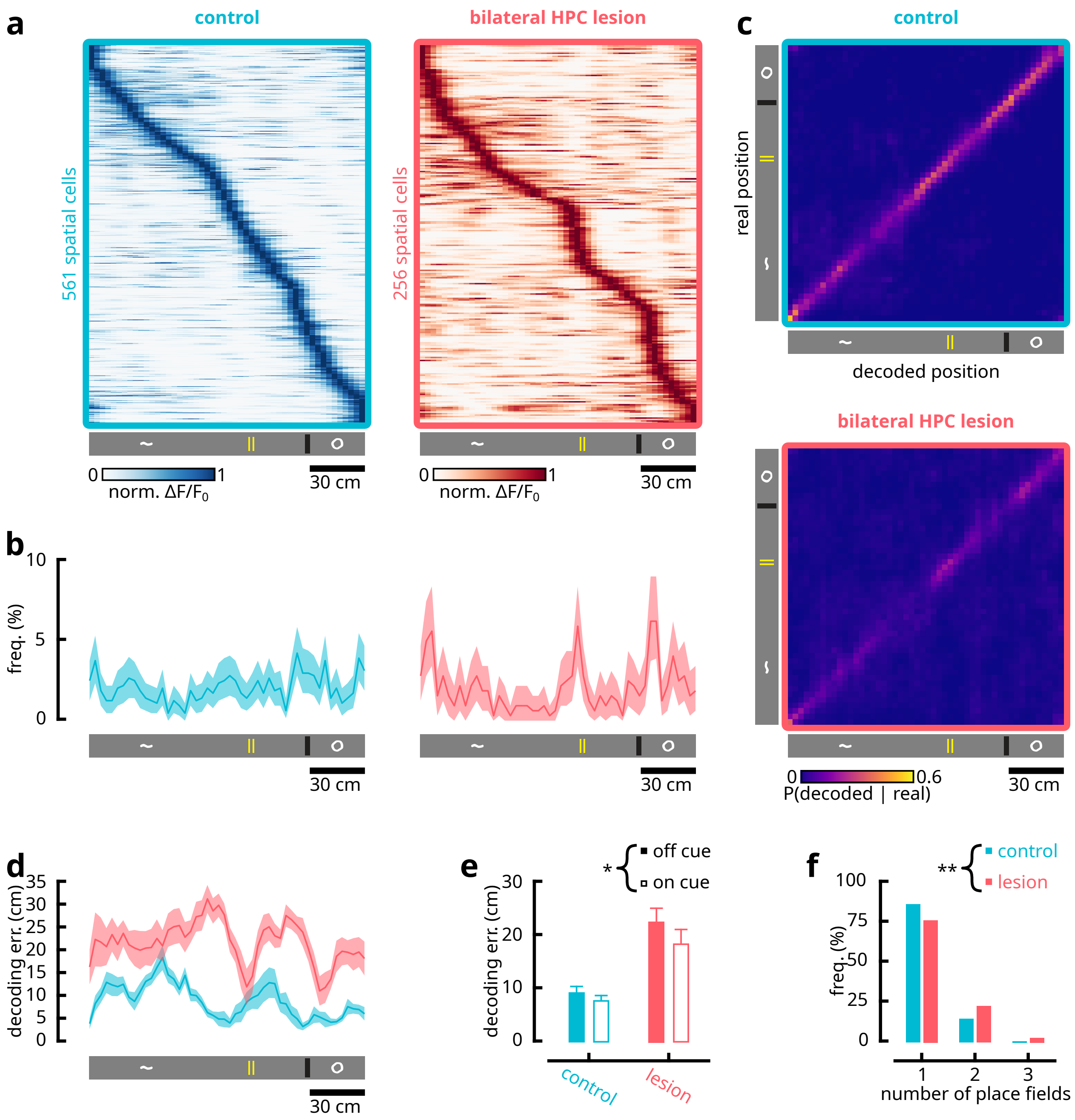

**Supplementary figure 3**

Representation of visuo-tactile cues is preserved in the secondary motor cortex following bilateral lesion of the dorsal hippocampus. Data from Esteves et al. (2021). (**a**) The average response (normalized between 0 to 1) of spatially-selective neurons as a function of spatial location for control (blue) and hippocampal-lesioned (red) animals. Neurons sorted by their peak average firing location. Note that the population representation of space in control animals is approximately uniform, while this representation is biased towards the locations of cues in the lesioned group. (**b**) Histogram distribution of place field centres in control (blue; $n = 650$ place fields) and lesioned (red; $n = 327$ place fields) groups. Shaded areas represent 95 % bootstrapped confidence intervals. (**c**) Normalized confusion matrices for real and Bayesian decoded positions (see Esteves et al. (2021) for methods). Decoding error assessed by *leave-one-out* cross-validation over trials. Notice that accuracy is higher over cue locations in the lesion group. (**d**) Average decoding as a function of spatial location ($n = 10$ sessions in $n = 4$ control animals; $n = 8$ sessions in $n = 4$ lesion animals). Shaded area denote S.E.M. (**e**) Average decoding error inside and outside cue locations. Robust two-way mixed ANOVA with 20 % trimmed-means (R package 'WRS'). Main effects of hippocampal lesion ($p = 0.010$) and cue location ($p = 0.028$). No significant interaction ($p = 0.080$). (**f**) Frequency in the number of place fields per individual spatial neurons. Spatial cells in the lesioned group tend to support a higher number of place fields ($\chi^2$ test; $p = 0.001$).

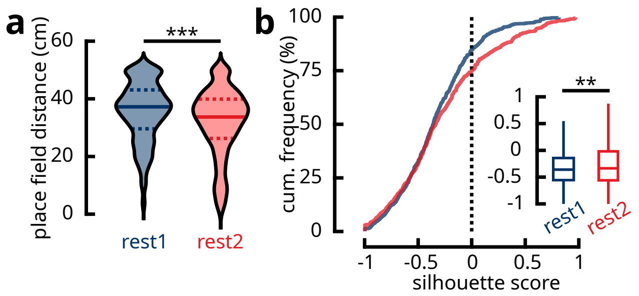

**Supplementary figure 4**

REST2 ensembles tend to be composed of spatially-selective neurons sharing neighbouring place fields. (**a**) The average distance between place field centres of neurons within the same REST1 (blue) or REST2 (red) ensemble ($n = 392$ REST1 ensembles; $n = 478$ REST2 ensembles; $p < 0.001$; Mann-Whitney U-test). The kernel densities for the violin plots were estimated using a 2 cm Gaussian window. The median (solid line), first and last quartiles (dashed lines) are shown. (**b**) Cumulative distribution functions of the silhouette coefficients of the place field centre locations of REST1 (blue) and REST2 (red) neurons that belonged to synchronous ensembles. The silhouette coefficient, in the current context, measures how similar the neurons within the same ensemble are as opposed to neurons outside the ensemble. Similarity is based on the Euclidean distance between place field centres. Silhouette values range from $-1$ to 1, where high values suggest strong cohesion between a neuron and the other neurons within the same ensemble, and clear separation of the neuron from to the rest of the neuronal population. A significantly larger portion of REST2 neurons contain positive silhouette values compared to REST1 neurons ($n = 2672$ REST1 ensemble neurons; $n = 3613$ REST2 ensemble neurons; $p < 0.001$; Kolmogorov–Smirnov test). Inset: silhouette coefficients rendered in boxplots ($p = 0.007$; Mann-Whitney U-test; outliers omitted).

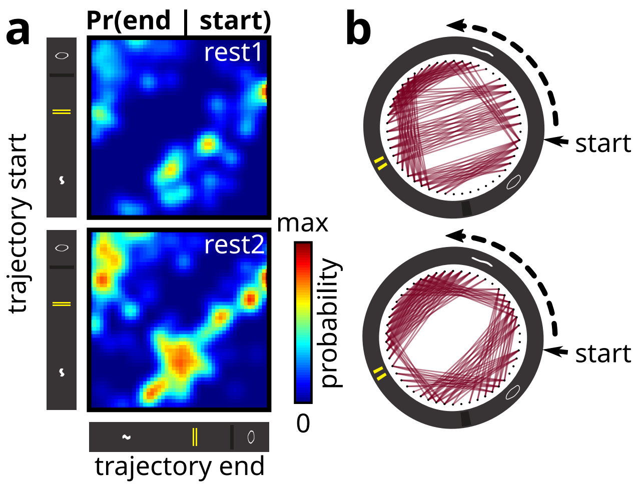

**Supplementary figure 5**

Trajectory ensembles encode short spatial segments that in many cases span the locations of cues. (**a**) Conditional probability matrices of the end location of trajectories given their starting location ($n = 160$ REST1 trajectories; $n = 325$ REST2 trajectories). Notice that the densities aggregate over a diagonal slightly offset from the central diagonal of the matrix, meaning that the majority the trajectories consist of short segments over space. Within these densities, a large fraction tend to span the locations of cues or the space delimited by two cues. Both of these trends are expressed prominently in REST2 trajectory ensembles, while REST1 ensembles show no clear organization. (**b**) The same probabilities in **b** represented in graph form. Short arrow point to the beginning of the track. Dashed arrow indicates the running direction.

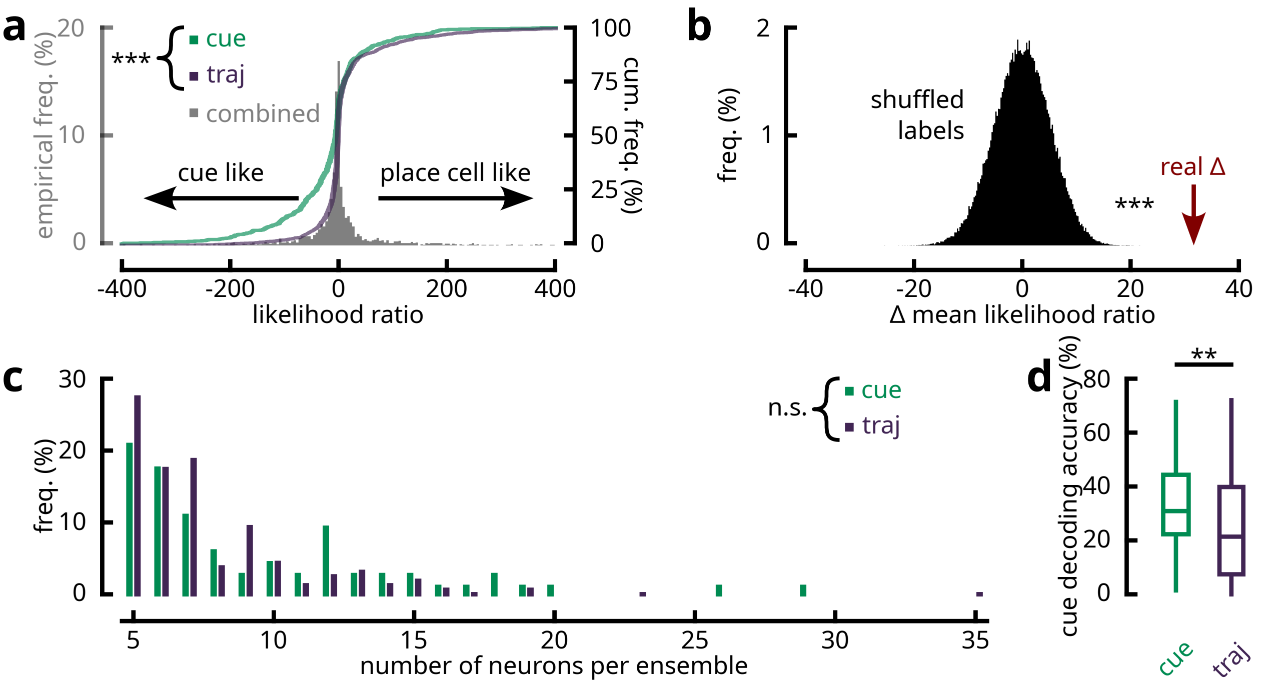

**Supplementary figure 6**

Cue and trajectory information are conjunctively encoded by resting-state ensembles; however, ensembles express varying degrees of bias for each separate behavioural feature. (**a**) Two models were fitted to the activities of ensemble neurons (see Methods): one is driven by sensory cues and the other is modelled after a *place-cell*'s tuning curve. Taking the likelihood ratio between the two models gives a neuron's proclivity for either type of response tuning. The distribution of these ratios express no bimodality, suggesting that ensemble neurons encode for a conjunction of cue and place responses. Labelling the neurons based on their cue/trajectory ensemble membership showed however that ensembles are biased in the type of behavioural features their encode for ($n = 601$ cue ensemble neurons; $n = 1314$ trajectory ensemble neurons; two-sample Kolmogorov–Smirnov test $p < 0.001$). (**b**) Distribution of the difference between mean likelihood ratios between cue and trajectory ensemble neurons, obtained by permutating the membership labels. The real difference is identified by the red arrow, confirming the bias for cue and trajectory ensembles in encoding their respective features ($p < 0.001$). (**c**) Distribution for the number of neurons per cue or trajectory ensemble. Cue and trajectory ensembles supported a similar number of neurons ($\chi^2$ test; $p = 0.17$), meaning that any detected differences in the encoding of behavioural parameters were not due to bias in sample size. (**d**) Bayesian decoding accuracy for the identity of individual cues using cue and trajectory ensemble neurons. Cue ensemble neurons were more accurate at identifying the identity of the four cues (two-tailed Mann-Whitney U-test; $p = 0.0012$), likely due to their tendency of supporting multiple place fields with different firing rates at different cue locations. However, this accuracy was still low at a median of $\sim$30 %. This could be due to the responses of cue ensemble neurons being targetted at specific cues rather than all four cues indiscriminately.

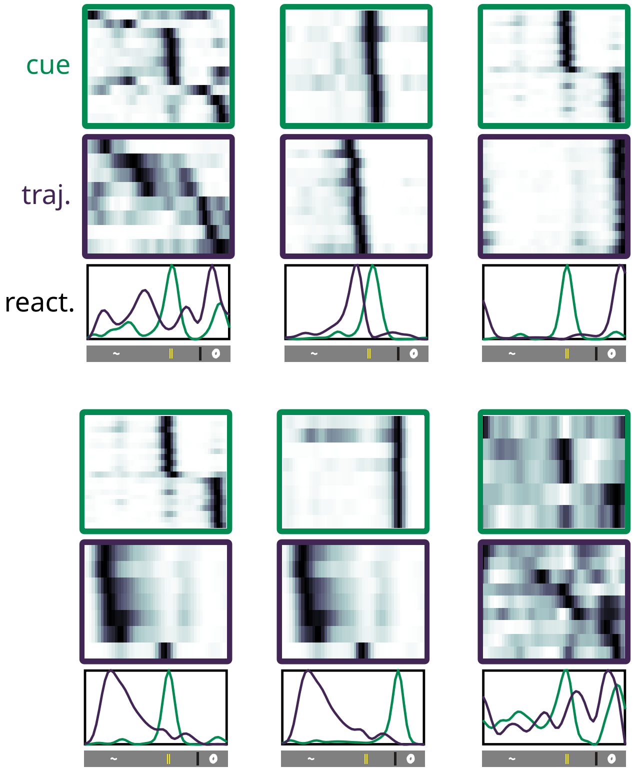

**Supplementary figure 7**

Six examples of temporally coupled cue-trajectory ensemble pairs. **top-middle:** Average neuronal activity as a function of spatial location for all neurons belonging to a cue or trajectory ensemble. Neurons were sorted by location of peak activity. **bottom:** The reactivation strength as a function of spatial location for the corresponding cue and trajectory ensembles.

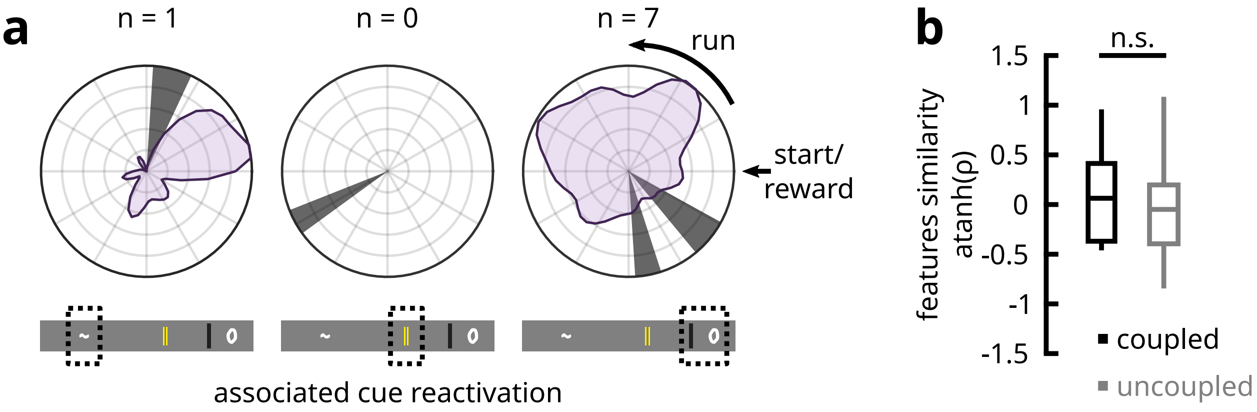

**Supplementary figure 8**

Cue and trajectory ensemble pairs from REST1 that are temporally-coupled. (**a**) Same as fig. 4**d** for coupled REST1 ensembles. Out of the eight pairs of cue and trajectory ensembles, seven were associated with the later two cues, while one is associated with the first cue. (**b**) Given the small sample size, the cue identify was omitted (cf. fig. 4**e**), and only the main effect of temporal-coupling was tested. No significant difference in the similarity of reactivated features between coupled and uncoupled ensemble pairs was detected (two-tailed two-sample t-test on atanh-transformed Pearson correlation coefficients; $p = 0.494$). However, given the low statistical power owing to the restricted sample size, it cannot be concluded that REST1 ensembles do not reactivate for complementary features.

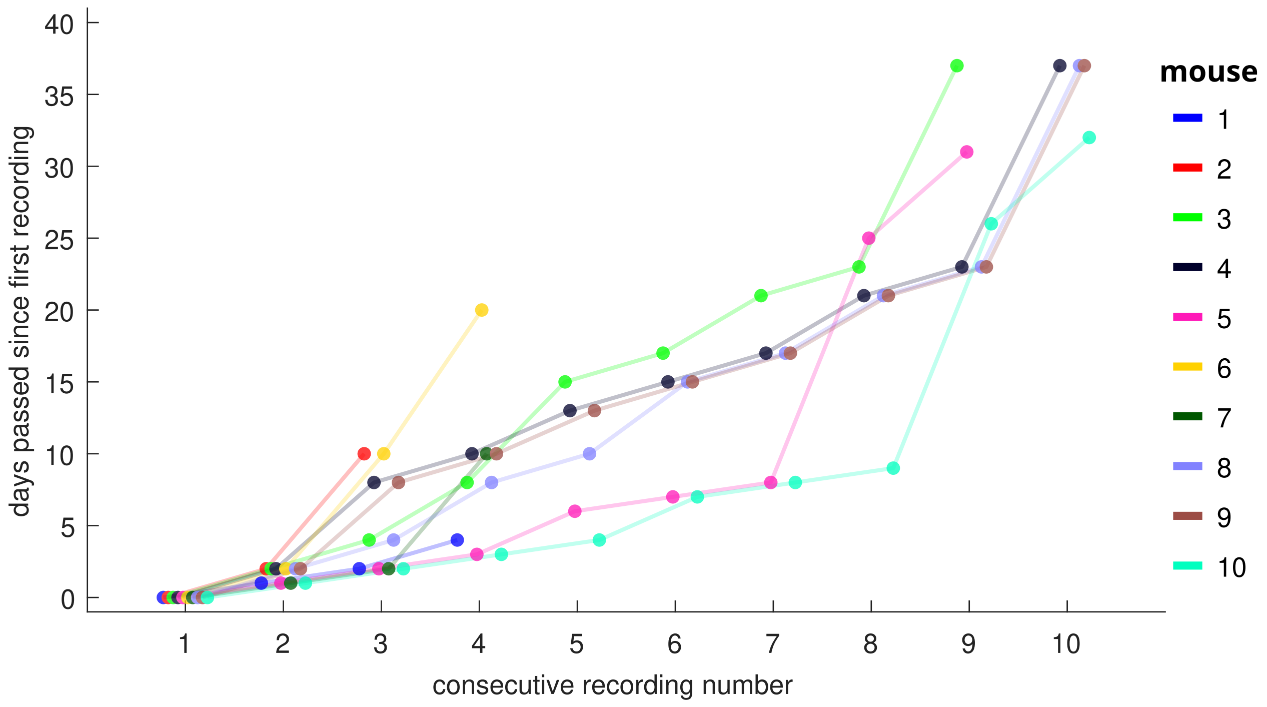

**Supplementary figure 9**

The number of days elapsed since the first recording for each consecutive recording sessions in the ten animals in which the same neuron ROIs were tracked across time.

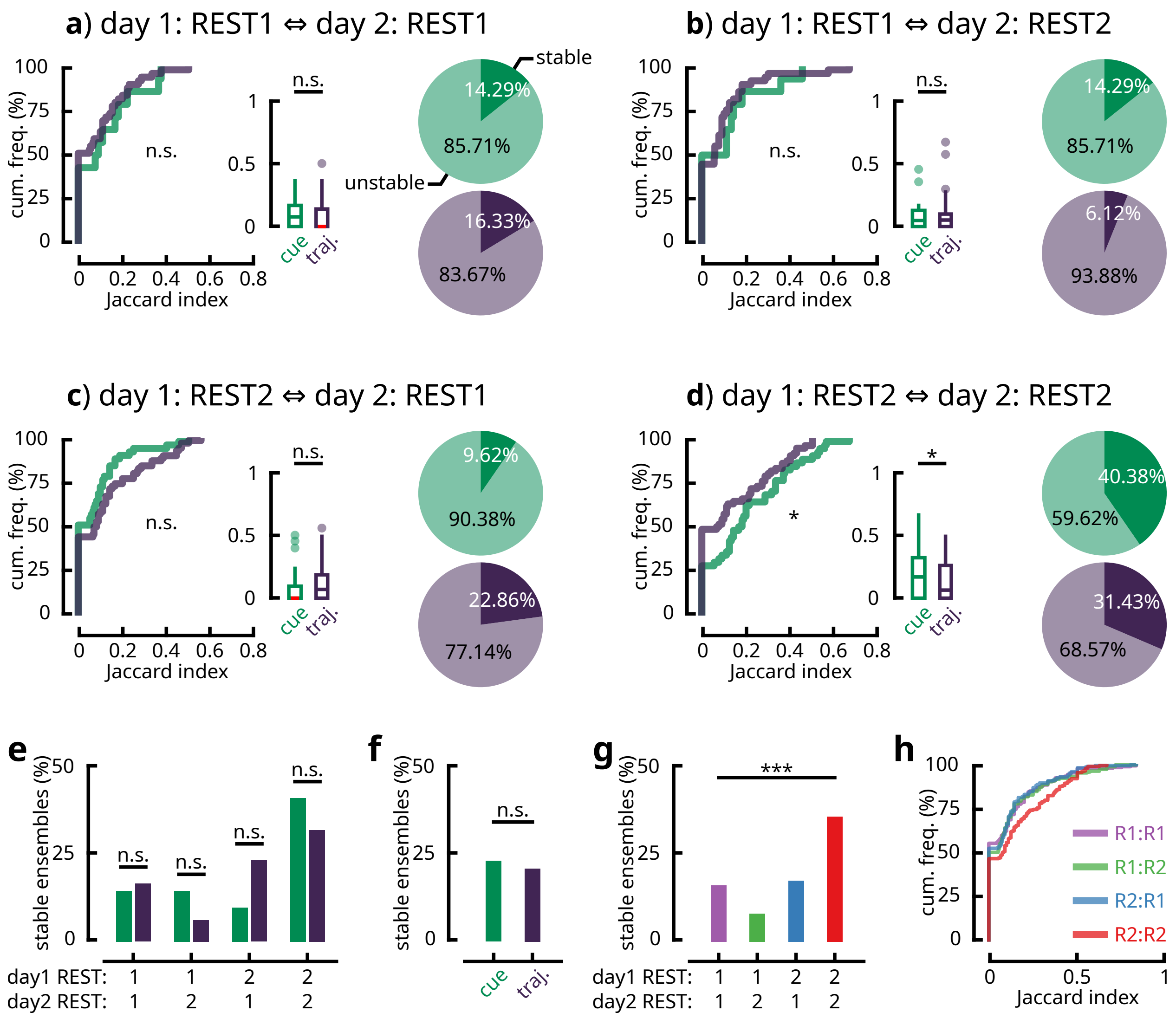

**Supplementary figure 10**

Resting state ensembles observed following active locomotion persist across recording days. (**a-d**) The proportions of overlapping ensemble members (quantified as Jaccard distances) and the percentage of persistent ensembles across consecutive recording days were determined using the same procedures described in fig. 5**e-f**. All combinations of REST1/REST2 ensembles across days were evaluated (e.g., in **a**, REST1 ensembles on the reference day were matched with REST1 ensembles on the subsequent recording day). Out of these combinations, cue ensembles ($n = 14$ in REST1; $n = 52$ in REST2) from REST2 on the reference day were slightly more stable than trajectory ensembles ($n = 49$ in REST1; $n = 70$ in REST2) in REST2 on the subsequent day (two-sample two-tailed Kolmogorov-Smirnov test $p = 0.024$; two-tailed Mann-Whitney U-test $p = 0.029$). Otherwise, no differences had been observed in the other combinations.

Overall, a greater fraction of both cue and trajectory ensembles persisted in REST2 across recording days (**d**). (**e**) To further corroborate the results in **a-d**, pairwise $\chi^2$ tests were performed between the proportions of persistent cue and trajectory ensembles over all REST combinations (df = 1; significance $\alpha = 0.05$). No differences between the fractions of stable cue and trajectory ensembles were found across all conditions, including in REST2-REST2, even prior to adjusting for multiple comparisons. Therefore, the increased persistence of cue ensembles across days following locomotion, observed in **d**, was marginal. (**f**) At the group level (i.e., taking the sum across all REST combinations), there was no difference in persistence between cue and trajectory ensembles either ($\chi^2 = 0.231$; df = 1; $p = 0.631$). (**g**) There was however a main effect of across-days REST combinations on the stability of ensembles, without accounting for cue/trajectory labels ($\chi^2 = 23.148$; df = 3; $p < 0.001$). (**h**) Plotting the empirical cumulative distribution functions of Jaccard distance for all REST combinations revealed that the proportions of stable cue or trajectory ensembles in REST2-REST2 was markedly higher than in other combinations. Taken together, these results suggest that following locomotion, similar resting state ensembles tend to be recruited across days, although no preference is assigned to either cue or trajectory ensembles.

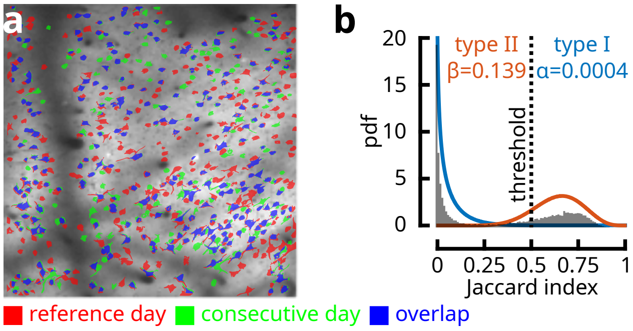

**Supplementary figure 11**

Identification of the same neuronal ROIs across consecutive days of recording. (**a**) Registered ROI masks for neurons detected on the reference day (red) and on the subsequent day of recording (green), in one example session. Overlapping pixels are coloured in blue. (**b**) Probability density function of the percentage of overlap (measured by Jaccard distance) between neuron ROIs across days, for all neuron pairs with at least one overlapping pixel. The histogram reveals a bimodal distribution consisting of persistent and differing cells. These two clusters were separated by K-means and were each fitted to a beta distribution by maximum likelihood estimation. The beta distribution, commonly used to model percentages and probabilities, is defined over the interval $[0, 1]$ and aptly describes Jaccard distances. With the overlapping threshold set at 50 %, the left-side beta distribution (blue), which represents unstable neurons, estimates a false positive rate of 0.04 %. Meanwhile, the statistical power given by the right-side distribution (orange) is estimated at 86.1 %. Therefore, a 50 % overlapping threshold for distinguishing persistent neurons across days is a highly robust criterion for the present dataset, at a reasonable cost to statistical power.

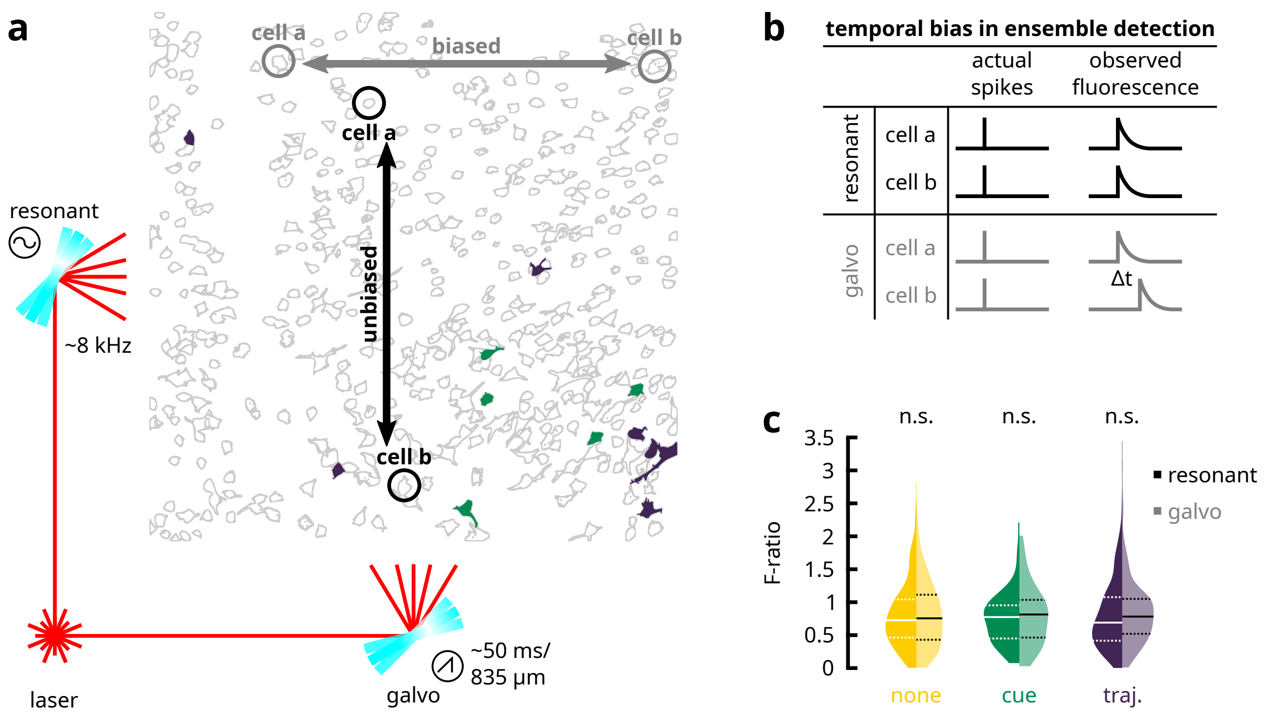

**Supplementary figure 12**

Detection of offline ensembles is not biased by the scan mirrors. (**a**) The laser beam is directed to the two axes of the field-of-view by a galvanometer scan mirror and a resonant scan mirror respectively. Examples of two offline ensembles (a cue ensemble in green and a trajectory ensemble in purple) are superimposed over all detected ROIs. (**b**) A direct consequence of this preparation is that neurons that are farther apart over the galvo axis (up to the midway point between the scanning path and the mirror flyback) are susceptible to shifted/delayed temporal dynamics. This can introduce a bias in the detection of offline ensembles, where synchronous ensembles found along the resonant axis are more likely to be detected compared to synchronous neurons along the galvo axis. Temporal smoothing had been performed to compensate for this issue. (**c**) To verify that this bias did not impact the detection, the following statistical hypothesis was proposed: the variance in the positions of ensemble neurons along the galvo axis, normalised by the total variance of all ROIs (i.e., the F-ratio), should be equal to that along the resonant axis. This null hypothesis held for all detected offline ensembles, irrespective of category (paired-sample two-tailed Wilcoxon signed rank tests; $p = 0.5$ for none; $p = 0.36$ for cue; $p = 0.273$ for trajectory). Therefore, synchronous neuron pairs are as likely to be detected over the span of the resonant axis as they are over the galvo axis.

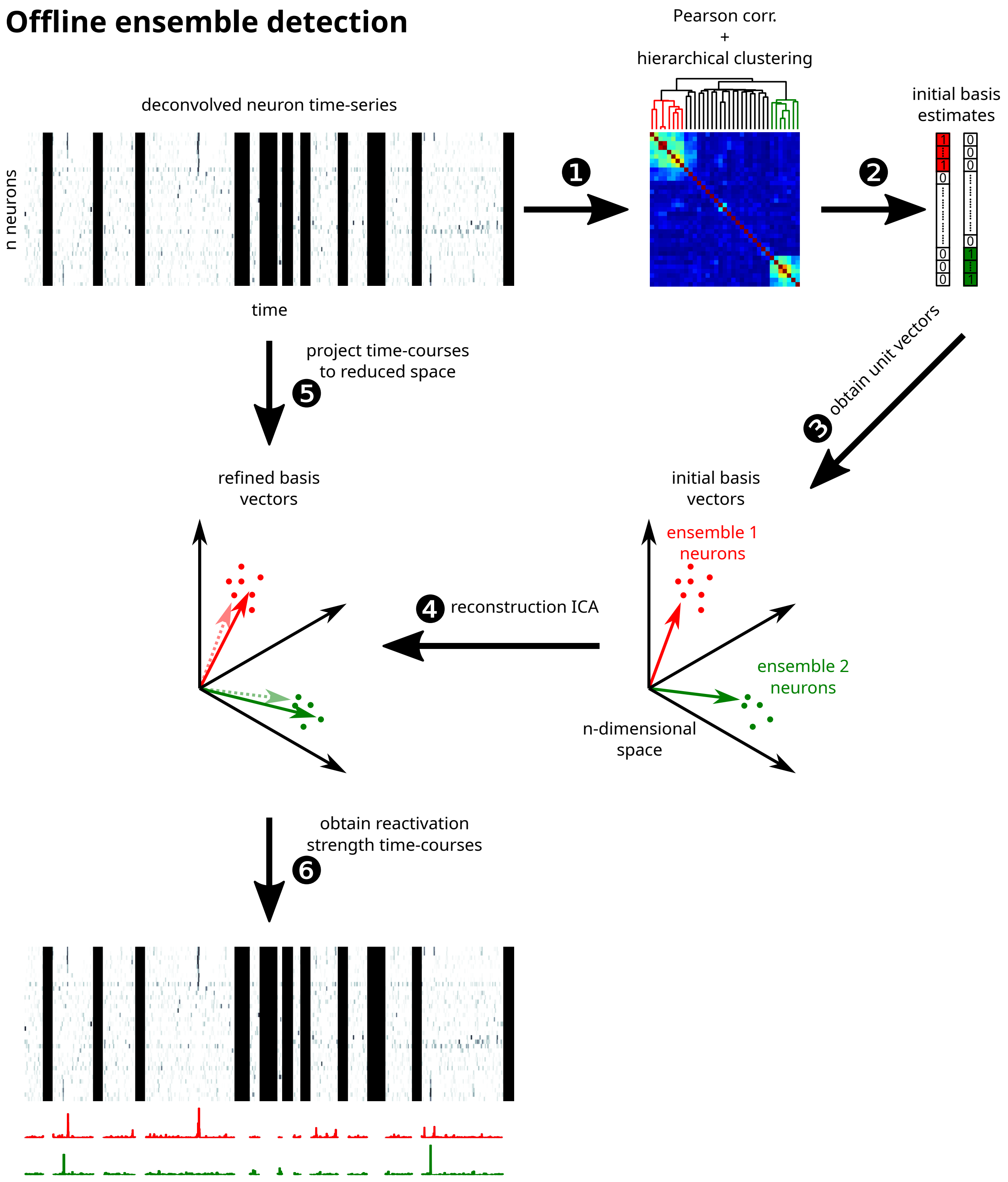

**Supplementary figure 13**

Methods used for detecting synchronous resting-state ensembles and for establishing the time-courses of their reactivation (see Methods for detailed description). 1) Hierarchical clustering is conducted over the Pearson correlation matrix of the neuronal time-series for the resting period. 2) For each ensemble, a binary column vector of the same length as the total number of neurons is created. Members belonging to the ensemble are labelled as ‘1’, while the remaining are ‘0’. 3) Normalizing these vectors by their norm yields a set of unit vectors that together form an orthonormal basis. 4) Using reconstruction ICA, these basis vectors are fine-tuned in such a way as to retain most of the variance in the original data, and in doing so capture the relative contributions/weights of each member neuron to an ensemble’s temporal dynamics. 5) The original neuronal time-series matrix is projected into the new space defined by the basis. 6) This projection yields the reactivation strength of each ensemble as a function of time.

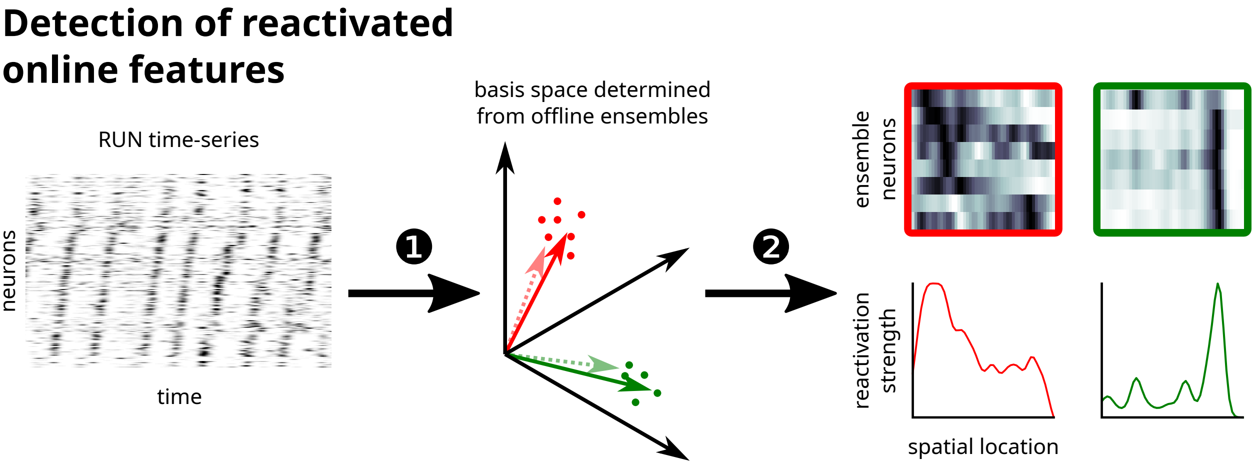

**Supplementary figure 14**

By projecting the neuronal time-series matrix during RUN into the space defined by the basis estimated from resting network dynamics (1), the ensemble activation strength as a function of animal location can be obtained (2).

| | | | | # ensembles | | | | | | | |
|---|---|---|---|---|---|---|---|---|---|---|---|
| | | | | REST1 | | | | REST2 | | | |
| **animal** | **# sessions** | **LFP** | **ROI tracking** | **none** | **cue** | **traj.** | **all** | **none** | **cue** | **traj.** | **all** |
| 1 | 6 | n/a | n/a | 12 | 0 | 7 | 19 | 13 | 0 | 10 | 23 |
| 2 | 5 | n/a | n/a | 24 | 2 | 10 | 36 | 16 | 3 | 34 | 53 |
| 3 | 1 | n/a | n/a | 6 | 1 | 2 | 9 | 0 | 0 | 2 | 2 |
| 4 | 4 | avail. | avail. | 10 | 0 | 3 | 13 | 9 | 1 | 2 | 12 |
| 5 | 3 | poor | avail. | 19 | 1 | 13 | 33 | 16 | 8 | 13 | 37 |
| 6 | 1 | avail. | n/a | 4 | 0 | 1 | 5 | 6 | 1 | 3 | 10 |
| 7 | 9 | avail. | avail. | 19 | 1 | 4 | 24 | 38 | 1 | 15 | 54 |
| 8 | 10 | avail. | avail. | 30 | 5 | 9 | 44 | 12 | 17 | 12 | 41 |
| 9 | 9 | avail. | avail. | 26 | 0 | 4 | 30 | 13 | 4 | 3 | 20 |
| 10 | 4 | avail. | avail. | 14 | 3 | 8 | 25 | 26 | 6 | 16 | 48 |
| 11 | 4 | poor | avail. | 9 | 1 | 15 | 25 | 7 | 10 | 17 | 34 |
| 12 | 10 | avail. | avail. | 15 | 1 | 0 | 16 | 12 | 1 | 8 | 21 |
| 13 | 10 | avail. | avail. | 50 | 1 | 7 | 58 | 52 | 0 | 10 | 62 |
| 14 | 10 | avail. | avail. | 38 | 3 | 14 | 55 | 32 | 10 | 19 | 61 |
| **total** | **86** | **9** | **10** | **276** | **19** | **97** | **392** | **252** | **62** | **164** | **478** |

**Supplementary table 1**

Experimental conditions for individual animal subjects.
